# Coordinated activity of sleep and arousal neurons for stabilizing sleep/wake states in *Drosophila*

**DOI:** 10.1101/243444

**Authors:** Jinfei D. Ni, Tyler H. Ogunmowo, Hannah Hackbart, Ahmed Elsheikh, Adishthi S. Gurav, Andrew A. Verdegaal, Craig Montell

## Abstract

The output arm of the sleep homeostat in *Drosophila* is a group of neurons with projections to the dorsal fan-shaped body (dFSB) of the central complex in the brain. However, neurons that regulate the sleep homeostat remain poorly understood. Using neurogenetic approaches combined with *ex vivo* Ca^2+^ imaging, we identify two groups of sleep-regulatory neurons that modulate the activity of the sleep homeostat in an opposing fashion. The sleep-promoting neurons activate the sleep homeostat with glutamate, whereas the arousal-promoting neurons down-regulate the sleep homeostat’s output with dopamine. Co-activating these two inputs leads to frequent shifts between sleep and wake states. We also show that dFSB sleep homeostat neurons release the neurotransmitter GABA that inhibits octopaminergic arousal neurons. Taken together, we suggest coordinated neuronal activity of sleep- and arousal-promoting neurons is essential for stabilizing sleep/wake states.

**Highlights:**

- Glutamate released by AstA neurons activates dFSB^AstAR1^ sleep-promoting neurons
- Dopamine down-regulates the activity of dFSB^AstAR1^ neurons
- Simultaneous glutamate and dopamine input causes rapid sleep and awake swings
- GABA released by dFSB^AstAR1^ neurons promotes sleep by inhibiting arousal neurons

## Introduction

Sleep is a fundamental physiological process conserved across a diverse range of animals. Most animals spend a large proportion of their life sleeping (Sehgal and Mignot, 2011). This promotes survival since deficits in sleep are linked to neurological diseases such as Alzheimer’s, Parkinson’s, depression, schizophrenia, as well as metabolic disorders and immunological dysfunction (Iranzo, 2016; Tsuneki et al., 2016; Westermann et al., 2015).

Sleep regulatory mechanisms that have been deciphered in model organisms such as flies and mice appear to be conserved in humans (Borbely, 1982; Griffith, 2013; Harbison et al., 2009; Hendricks et al., 2000; Sehgal and Mignot, 2011; Shaw et al., 2000; Tomita et al., 2017; Weber and Dan, 2016). These include dual regulation by the circadian clock and by homeostatic-drive, both of which are essential for maintaining appropriate sleep patterns (Allada et al., 2017; Borbely, 1982). For instance, increased sleep drive due to sleep deprivation leads to compensatory sleep. In *Drosophila*, a group of neurons project to the dorsal fan-shaped body (dFSB neurons) of the central complex in the brain, and are proposed to be the effector component of the sleep homeostat (Donlea et al., 2014; Liu et al., 2016).

Sleep is essential for animal survival, but comes with a large trade-off. Other critical behaviors, such as feeding, mating and defense, only take place when an animal is awake (Griffith, 2013). Therefore, arousal-promoting neurons that stimulate wakefulness are essential. In *Drosophila*, various arousal signals have been identified and their functions in promoting wakefulness are conserved in mammals (Crocker et al., 2010; Liu et al., 2012; Sehgal and Mignot, 2011; Ueno et al., 2012a). These include biogenic amines (dopamine and octopamine) and neuropeptides such as Pigment Dispersing Factor (PDF) (Parisky et al., 2008) (Andretic et al., 2005; Crocker and Sehgal, 2008; Ueno et al., 2012b). Animals transit between wake and sleep states by rapid changes in a host of physiological processes including sensory processing, motor output, memory consolidation, and metabolism. However, the coordination of sleep-promoting and wake-promoting neuronal circuits is poorly understood.

To address the fundamental question as to how antagonistic sleep and arousal pathways are coupled, we combined genetics, neuroanatomical and neurophysiological approaches in *Drosophila*. We found that dFSB neurons, which express the Allatostatin receptor1, AstAR1 (dFSB^AstAR1^), are activated by sleep-promoting neurons that release glutamate and express the neuropeptide, Allatostatin-A (AstA). We also find that dopaminergic wake-promoting neurons form synaptic connections at both dendritic and axonal projections of the dFSB^AstAR1^ sleep-promoting neurons. Dopamine released by arousal (DAA) neurons down-regulates the activity of dFSB^AstAR1^ neurons and inhibits their synaptic output at the axonal terminal. When excitatory glutamate and inhibitory DA input impinge on dFSB^AstAR1^ neurons simultaneously, the animals exhibit frequent transitions between sleep and wake states. We also found that dFSB^AstAR1^ neurons promote sleep by releasing GABA which inhibits previously identified octopaminergic arousal (OAA) neurons (Crocker et al., 2010), thereby conferring long sleep bouts at night. Our work provides a molecular and neuronal circuit mechanism as to how sleep-promoting neurons communicate with arousal-promoting neurons to coordinate sleep.

## Results

### Activation of Allatostatin-A neurons promotes sleep

To dissect the neuronal circuits that regulate sleep in *Drosophila*, we expressed the warm-activated TRPA1 channel (*UAS-trpA1*) in different subsets of putative peptidergic neurons using a variety of *Gal4* drivers. TRPA1 is activated at 29° but not 22°C (Parisky et al., 2008; Viswanath et al., 2003). Therefore, if we drive TRPA1 expression in sleep-promoting neurons and increase the ambient temperature to 29°C, the animals would display an increase in sleep relative to 22°C. On the other hand, if a *Gal4* line drives *UAS-trpA1* expression in neurons that promote arousal, total sleep would decline at 29°C.

We identified three independent *Gal4* lines that promote sleep upon thermal activation by TRPA1 (Figure S1; 39351, 51978, 51979). All three lines were expressed under the transcriptional control of the gene encoding the neuropeptide Allatostatin-A (AstA) (Hergarden et al., 2012). Given the similar effects of these lines, we focused on one *AstA-Gal4* line (39351) to conduct the following investigation. Control flies (*UAS-trpA1/+* only, or *AstA-Gal4/+* only) exhibit only small increases in total sleep time after shifting the temperature from 22°C to 29°C (Figures 1A, 1B and 1D). In contrast, activating *AstA-Gal4* neurons in *AstA>trpA1* flies by increasing the temperature to 29°C drastically increased the amount of total sleep (Figures 1C and 1D), as previously reported (Chen et al., 2016).

**Figure 1.**
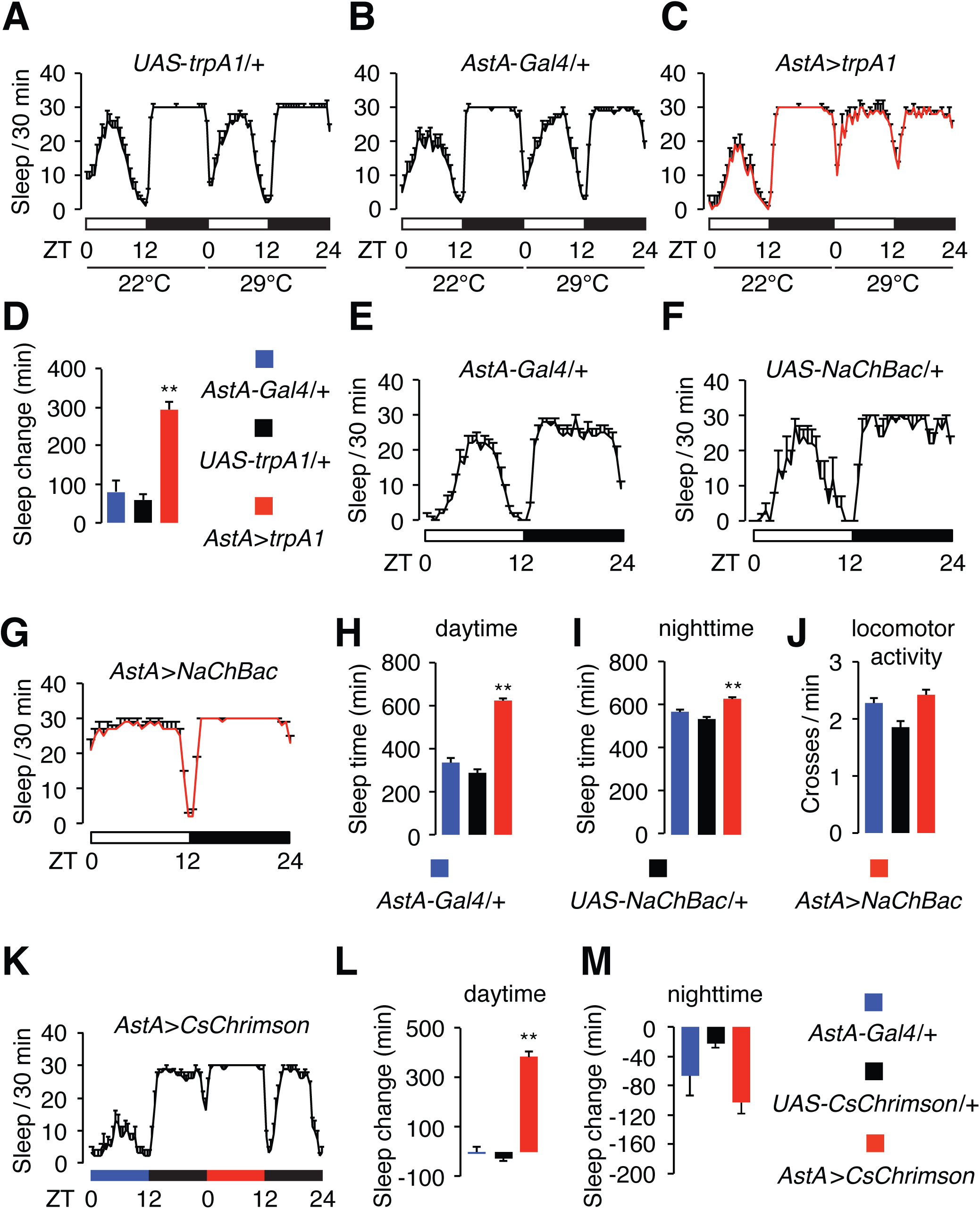
Activation *AstA-Gal4* neurons promotes sleep. (A—C) Sleep profiles of control flies (*UAS-trpA1/+* and *AstA-Gal4/+*) and flies expressing *UAS-trpA1* using the *AstA-Gal4* (*AstA>trpA1*). The white and black bars beneath the sleep profiles indicate the day and night cycles, respectively. ZT0 and ZT12 indicate the times when lights are turned on and off, respectively. The experimental temperatures (22° and 29°C) are indicated below the white and back bars. The sleep times are plotted in 30-min bins. (D) Quantification of the increase in sleep over the course of 24 hours, resulting from thermal activation of TRPA1 at 29°C (Total sleep over the course of 24 hours at 29°C minus total sleep over the course of 24 hours at 22°C). n=23−78. (E—G) Sleep profiles of control flies (*AstA-Gal4/+* and *UAS-NaChBac/+*) and *AstA>NaChBac* flies. (H — I) Quantification of the daytime and nighttime sleep exhibited by the indicated flies. n=16−68. (J) Quantification of locomotor activity of the indicated flies during wakefulness. n=16−68. (K) Red light activation of CsChrimson in *AstA-Gal4* neurons increases sleep. Red but not blue light activates CsChrimson. Flies were entrained under 12-hour blue light/dark cycles for 3 days, and then shifted to a red light/dark cycle on the 4^th^ day. Shown is the sleep profile under the blue and red light conditions, as indicated by the blue and red bars. (L—M) Quantification of daytime and nighttime sleep change induced by activation of CsChrimson. *AstA-Gal4/+* and *UAS-CsChrimson/+* served as non-activated controls. n = 32–70. Error bars, SEMs. **p<0.01, one-way ANOVA with Dunnett’s test.

To assess the effects of depolarizing *AstA-Gal4* neurons in the absence of a temperature change, we used two approaches. First, we expressed NaChBac, a bacterial Na+ channel (Nitabach et al., 2006) in *AstA-Gal4* neurons. Compared to the controls (*AstA-Gal4/+* only, and *UAS-NaChBac/+* only), flies that expressed *UAS-NaChBac* under control of the *AstA-Gal4* (*AstA>NaChBac* flies) showed a substantial increase in sleep, which was most profound during the day (Figures 1E—1H). Nighttime sleep was also enhanced, although the change was less pronounced since flies sleep extensively at night (Figures 1E—1G and 1I). Depolarizing *AstA-Gal4* neurons did not appear to impair the flies’ locomotive ability because *AstA>NachBac* flies were as active as the control flies during the wake periods (Figure 1J).

In a second temperature-independent approach, we expressed a red-light activated channel-rhodopsin, CsChrimson (Klapoetke et al., 2014) in *AstA-Gal4* neurons. We entrained flies using a 12-hour blue-light/12-hour dark paradigm, which did not activate CsChrimson, and then employed a red-light/dark cycle. We observed a large increase in daytime sleep upon neuronal activation with red light (Figures 1K and 1L). Following the daytime activation of *AstA-Gal4* neurons, the amount of sleep during the following night declined, although the difference from the control was not statistically significant (Figures 1M).

### Two groups of sleep-promoting neurons labeled by the *AstA-Gal4:* SLP^AstA^ and LPN^AstA^

To identify the sleep-promoting neurons labeled by the *AstA-Gal4*, we first characterized the expression pattern of the *AstA-Gal4* using a membrane-tagged GFP (*UAS-mCD8::GFP*) and performed whole-mount immunofluorescence of the fly brain. GFP labeled several regions in the *Drosophila* brain, including neurons that innervate the medulla layer of the optical lobe (med), neurons innervating the primary gustatory center, the subesophageal zone (SEZ), neurons that project to the superior lateral protocerebrum (SLP), and three neurons located at lateral posterior region of the brain (LPN), which are circadian pacemaker neurons (Figures 2A and 2B). We confirmed the identity of the LPN neurons by co-staining with Timeless (Tim; Figures 2C, 2D and S2A), one of the core clock proteins expressed in the circadian pacemaker neurons (Myers et al., 1996). These LPNs also stained with anti-AstA, and are referred to here as LPN^AstA^ neurons (Figures 2E and 2F). We generated a knock-in reporter (*AstA^LexA^*) by replacing the *AstA* coding region with *LexA* using CRISPR-mediated homologous recombination (Bassett et al., 2013; Kondo and Ueda, 2013; Ren et al., 2013). We then expressed mCD8::GFP using this *AstA^LexA^* reporter and found that it also labeled the three LPN^AstA^ neurons, but not SLP^AstA^ neurons (Figure 2G).

**Figure 2.**
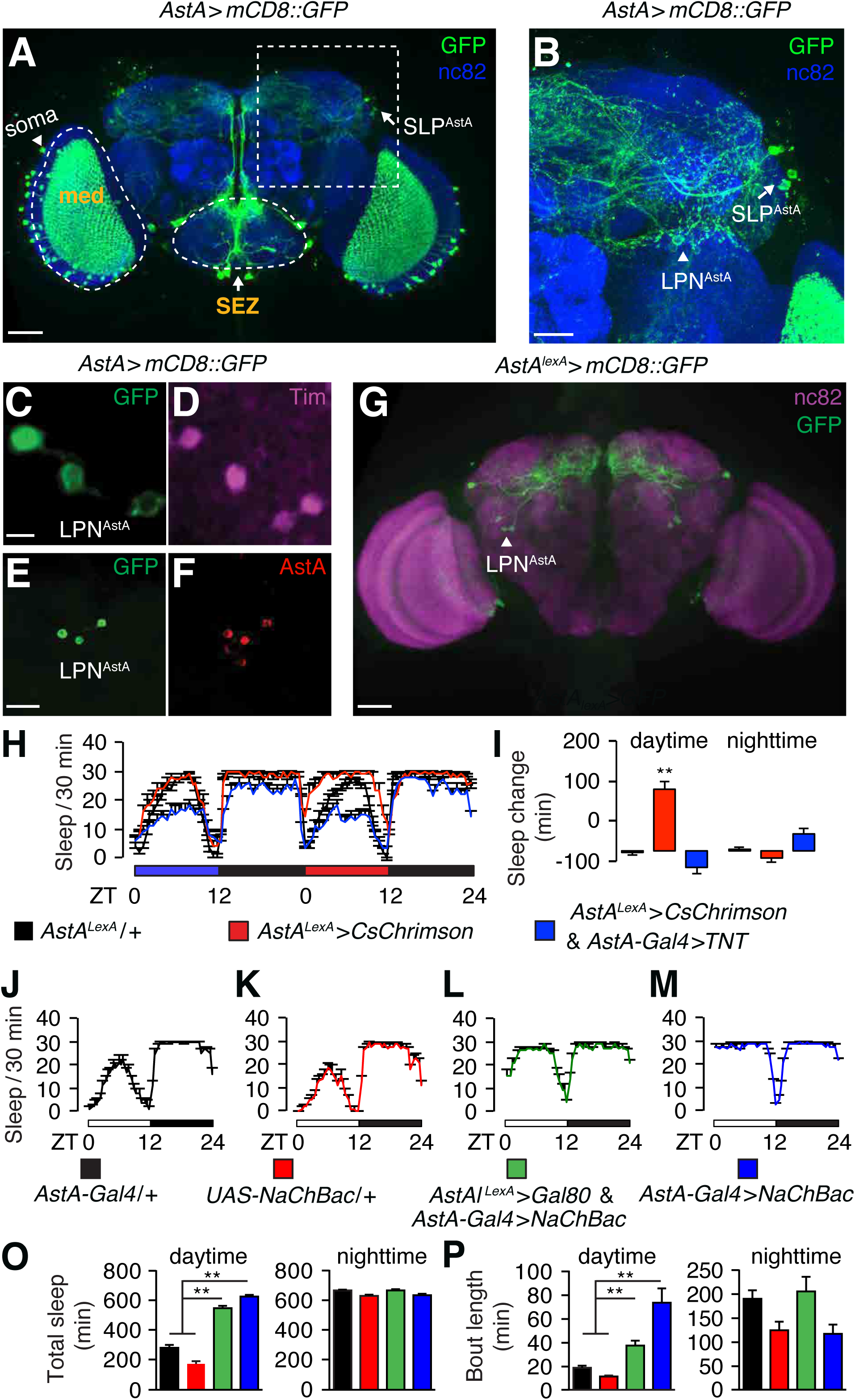
Identification of a subset of neurons labeled by *AstA-Gal4* as the sleep-promoting neurons. (A) Immunostaining of a whole-mount of a brain from a fly expressing *UAS-mCD8::GPF*under the control of the *AstA-Gal4* (*AstA>mCD8::GFP*). Green, anti-GFP; blue: anti-NC82. The dashed box indicates the region shown at higher magnification in B. The scale bar represents 40 μm. (B) Zoom-in view of boxed region in A. The arrow and arrowhead indicates SLP^AstA^ and LPN^AstA^ neurons, respectively. The scale bar represents 20 *μ*m. (C—F) Immunostaining of LPN^AstA^ neurons in brain whole-mounts from *AstA>mCD8::GFP* flies. C and D show co-staining with anti-GFP (green) and anti-Tim (magenta). The scale bar represents 5 *μ*m. E and F show co-staining of anti-GFP (green) and anti-AstA (red). The scale bar represents 20 *μ*m. (G) Immunostaining of a brain whole-mount (*AstA^LexA^> lexAop-mCD8::GFP*) with anti-GFP (green) and anti-NC82 (magenta). The scale bar represents 40 *μ*m. (H) Sleep time (plotted in 30-min bins) of the indicated flies exposed to blue light (non-activation of CsChrimson) or red light (activation of CsChrimson). (I) Quantification of the changes in daytime and nighttime sleep by red light induced activation of CsChrimson. n=10−32. Note that the red light induced sleep-promoting effect of *AstA^LexA^>CsChrimson* is abolished when in combination with *AstA-Gal4>UAS-TNT*. (J — K) Sleep profiles of: (J) *AstA-Gal4/+*, (K) *UAS-NaChBac/+*, (L) *AstA^LexA^>Gal80* and *AstA>NaChBac*, and (*M*)*AstA>NaChBac*. (O) Quantification of daytime sleep and nighttime sleep of the flies shown in J — K. (P) Quantification of daytime sleep and nighttime sleep-bout lengths of the flies shown in J — K. n=23−32. Error bars, SEMs. **p<0.01, one-way ANOVA with Dunnett’s test.

To identify which subsets of the *AstA-Gal4* neurons are sleep promoting, we used a genetic “*FlpOut*” approach to generate flies expressing TRPA1 tagged with mCherry (mCherry::TRPA1) in different subgroups of neurons labeled by the *AstA-Gal4* (Figure S3A and S3B). To induce stochastic expression of mCherry::TRPA1 in subsets of *AstA-Gal4* neurons, we heat-shocked flies during the 3^rd^ instar larval period to induce expression of the Flipase (Flp), which would remove the stop codon flanked by FRT sites (Figures S3A and S3B). After eclosion, we monitored the sleep of individual *AstA>FlpOut-mCherry::TRPA1* flies and plotted the changes in sleep exhibited by individual flies upon thermal activation of mCherry::TRPA1 (Figure S3C). Without inducing Flp expression with heat shock, flies exhibited a similar level of sleep regardless of whether the temperature was 22° or 29°C (Figure S3D).

To visualize the key sleep-promoting neurons, we compared the expression patterns of *mCherry::trpA1* in the brains of *AstA>FlpOut-mCherry::trpA1* flies that did not show an increase in total sleep with the staining patterns from flies that showed significant increases in sleep upon thermoactivation. For comparison, we also expressed *UAS-GFP* under control of the *AstA-Gal4*. Neurons located in the medullar layer (med) of the optic lobe and the SEZ did not show differential expression between the two groups of flies, indicating they are not the sleep-promoting neurons (Figures S3E—S3H). In contrast, LPN^AstA^ neurons stained strongly with mCherry in flies that showed significant increases in sleep, and stained weakly in flies that did not show an increase in sleep (Figure S3I and S3J. Moreover, these LPN^AstA^ neurons, projected to the SMP (Figures S2 and S3I). We also found that mCherry stained SLP neurons (referred to as SLP^AstA^ neurons) that projected to the SMP only in flies manifesting an increase in sleep (Figures S2, S3I and S3J). These data indicate that LPN^AstA^ and SLP^AstA^ are sleep-promoting neurons.

To test whether activation of either LPN^AstA^ or SLPAstA neurons is sufficient to promote sleep, we specifically activated each of these neurons, and examined the impact on sleep. To activate LPN^AstA^ neurons, we expressed *CsChrimson* under control of the *AstA^LexA^* reporter, which labels LPN^AstA^, but not SLP^AstA^ neurons (Figures 2G). Activation of LPN^AstA^ neurons with red light led to an increase in daytime sleep, compared to control flies that did not express CsChrimson (Figures 2H and 2I). This sleep-promoting effect was diminished when we expressed tetanus toxin (TNT) using the *AstA-Gal4*, which labels LPN neurons, but not other neurons labeled by the *AstA^LexA^* reporter (Figures 2H—2I). These data indicate that neurotransmission from LPN^AstA^ neurons is required to promote sleep.

To test whether activation of SLP^AstA^ neurons is sufficient to promote sleep, we restricted *AstA-Gal4* expression to SLP^AstA^ neurons. To do so, we expressed the *Gal4* inhibitor, *Gal80* (Lee, 2009) under the control of the *AstA^LexA^*, thereby limiting *Gal4* expression to LPN^AstA^ neurons. We introduced *UAS-NaChBac* into this genetic background and found that these flies exhibited an increase in total daytime sleep and bout length relative to control flies (Figures 2O and 2P). However, the sleep-promoting effect was larger in flies expressing *UAS-NaChBac* in all *AstA-Gal4* neurons (Figures 2O and 2P). Nighttime sleep was not increased, presumably due to a ceiling effect from the high level of sleep in control flies (Figures 2O and 2P). We also observed a daytime sleep-promoting effect when we used CsChrimson to activate SLP^AstA^ neurons (Figures S4A).

To further test the contributions of SLP^AstA^ and LPN^AstA^ activation to sleep promotion, we combined the *AstA-Gal4* with the *tsh-Gal80*, which eliminated expression from these neurons (Figure S3K), and with two other *Gal80* lines (*ChaT-Gal80* and *Vglut-Gal80*) that did not impact on expression of SLP^AstA^ and LPN^AstA^ neurons (Figures S3L and S3M). We found that the sleep-promoting effects of either *UAS-trpA1* or *UAS-NaChBac* were either eliminated or greatly diminished using the *tsh-Gal80*, but were unaffected by introduction of either the *ChaT-Gal80* or the *Vglut-Gal80* (Figures S3N and S3O). These data support the proposal that SLP^AstA^ and LPN^AstA^ are the sleep-promoting neurons labeled by *AstA-Gal4*.

### *AstAR1-Gal4* labels sleep-promoting neurons (dFSB^AstAR1^) that project to the dFSB

To identify sleep-promoting neurons that could potentially function post-synaptic to the neurons labeled by *AstA-Gal4*, we interrogated neurons that were labeled by reporters for the receptor for AstA (AstAR1). Therefore, we expressed *UAS-NaChBac* under the control of a collection of *AstAR1 Gal4s* (see Method Details and Jenett et al., 2012). Among the different *AstAR1-Gal4* lines, 23E10 had the strongest effect, and resembled the sleep-promoting phenotype of *AstA>NaChBac* flies during the day (Figures 3A—3E). Because sleep was already high at night, there was little additional nighttime sleep in these flies (Figure 3F). This phenotype was not due to a general defect in locomotion since the overall activity of *AstAR1>NaChBac* flies during wake bouts was not decreased relative to control flies (Figure 3G). We expressed *UAS-CsChrimson* in *AstAR1-Gal4* neurons, and found that activating these neurons by shifting the light from blue to red drastically increased the daytime sleep relative to control flies (*AstAR1>CsChrimson/+;* Figures S4B—S4F).

**Figure 3.**
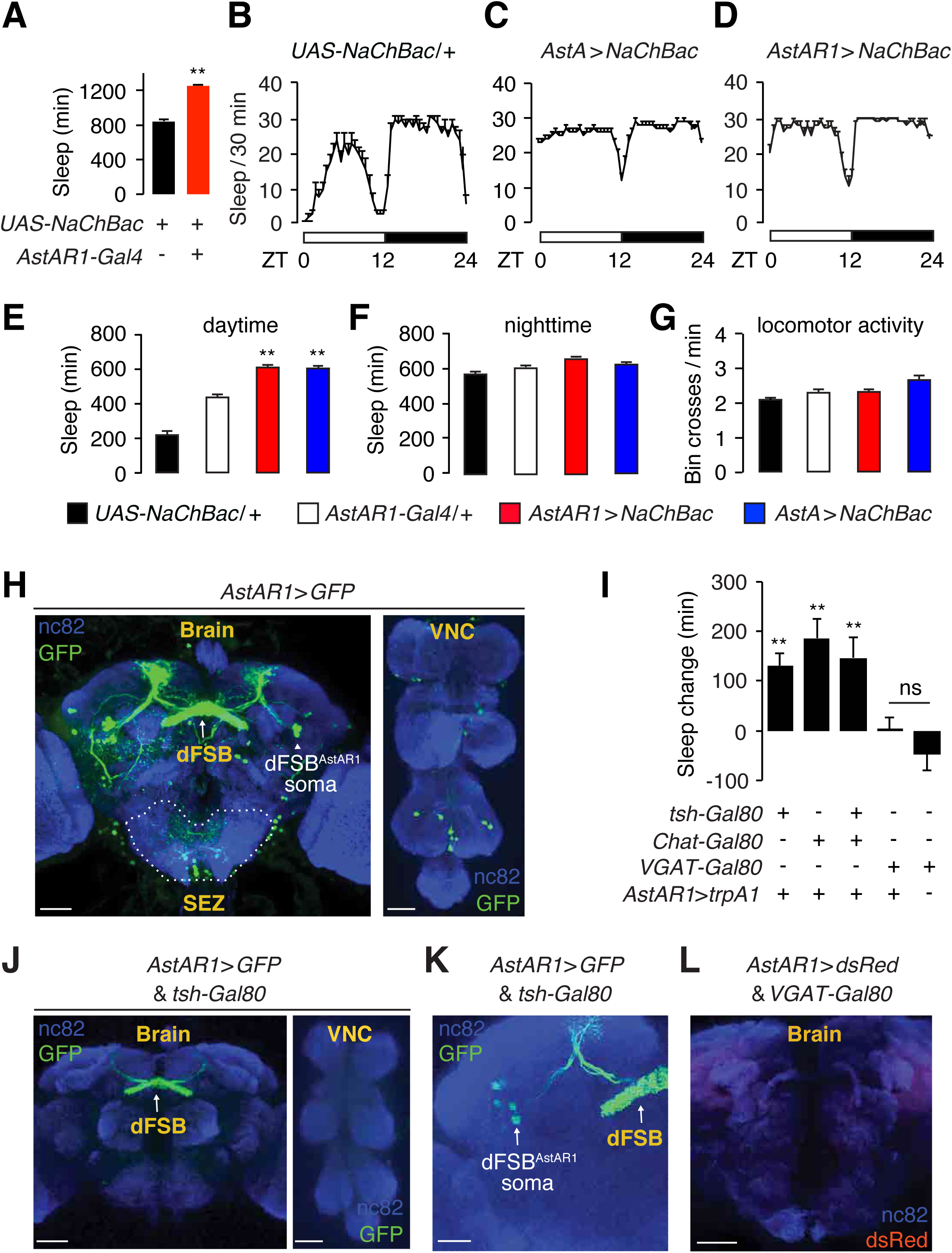
dFSB sleep-promoting neurons labeled by an *AstAR1-Gal4* reporter. (A) Effects on total sleep due to expression of *UAS-NaChBac* under control of the *AstAR1-Gal4*. Shown are the total minutes of sleep over a 24-hour light/dark cycle. n=16−53. (B—D) Sleep profiles of the indicated flies. (E—F) Quantification of daytime and nighttime sleep exhibited by the indicated flies. n=16−48. (G) Quantification of the locomotor activity of the indicated flies during periods of wakefulness over a 24-hour period. n=16−48. (H) Whole-mount brain and VNC of *AstAR1>GFP* flies (*AstAR1-Gal4* and *UAS-mCD8::GFP*) with anti-GFP (green), and anti-NC82 (blue). The scale bars represent 60 *μ*m. (I) Quantification of the change in total sleep over a 24-hour period due to activation of TRPA1 at 29°C. n=18−33. (J — K) Immunostaining of brains and VNC in *AstAR1>GFP* flies carrying the *tsh-Gal80* to restrict *AstAR1-Gal4* expression to the dFSB. Anti-GFP (green), and anti-NC82 (blue). The scale bars represent 60 *μ*m in J, and 30 *μ*m in K. (**L**) Immunostaining of a brain from a *AstAR1>dsRed* fly carrying the *VGAT-Gal80* to eliminate expression in the dFSB. The scale bar represents 40 *μ*m. Error bars, SEMs. **p<0.01. Unpaired Student’s t-test for panel A, and one-way ANOVA with Dunnett’s test for panels E—G and I.

We examined the expression pattern of the *AstAR1-Gal4* and found that it labeled neurons innervating the dFSB (Figure 3H), which is proposed to be one of the sleep-regulating centers (Donlea et al., 2014; Liu et al., 2016). The *AstAR1-Gal4* also sparsely labels neurons innervating the SEZ and the ventral nerve cord (VNC; Figure 3H)

To identify which groups of neurons labeled by the *AstAR1-Gal4* promote sleep, we applied a genetic intersection method by introducing different *Gal80s* into *AstAR1>trpA1* flies. The sleep-promoting effect exhibited by *AstAR1>trpA1* flies was preserved when we combined the *AstAR1-Gal4* with the *tsh-Gal80*, the *Chat-Gal80*, or both (Figure 3I), which eliminate most reporter staining except for the dFSB (Figures 3J and 3K and Figures S4G—S4I). The increased sleep was abolished by introduction of the *VGAT-Gal80* (Figure 3I), which eliminates expression in dFSB neurons (Figure 3L). We refer to these sleep-promoting neurons as dFSB^AstAR1^.

### dFSB^AstAR1^ form close associations with SLP^AstA^ and LPN^AstA^ neurons

Since activating dFSB^AstAR1^ neurons phenocopies the sleep-promoting effect of activation of SLP^AstA^ and LPN^AstA^ neurons, we tested whether or not the projections of these two groups of neurons form close associations, which would occur if they formed synapses. As a first approach, we imaged the relative dendritic and axonal locations of these neurons using a dendritic marker (DenMark) (Nicolai et al., 2010) and an axonal (syt::eGFP) (Zhang et al., 2002), which consists of eGFP fused to synaptotagmin—a vesicular synaptic protein located proximal to the Ca^2+^ microdomain at axonal terminals. Neurons labeled with the *AstA-Gal4* extend dendrites and axonal projections into the superior medial protocerebrum (SMP) region of the brain (Figures 4A and 4B), whereas neurons labeled with the *AstAR1-Gal4* send dendrites into the SMP, and axonal terminals primarily to the dFSB (Figures 4C, 4D and S2).

**Figure 4.**
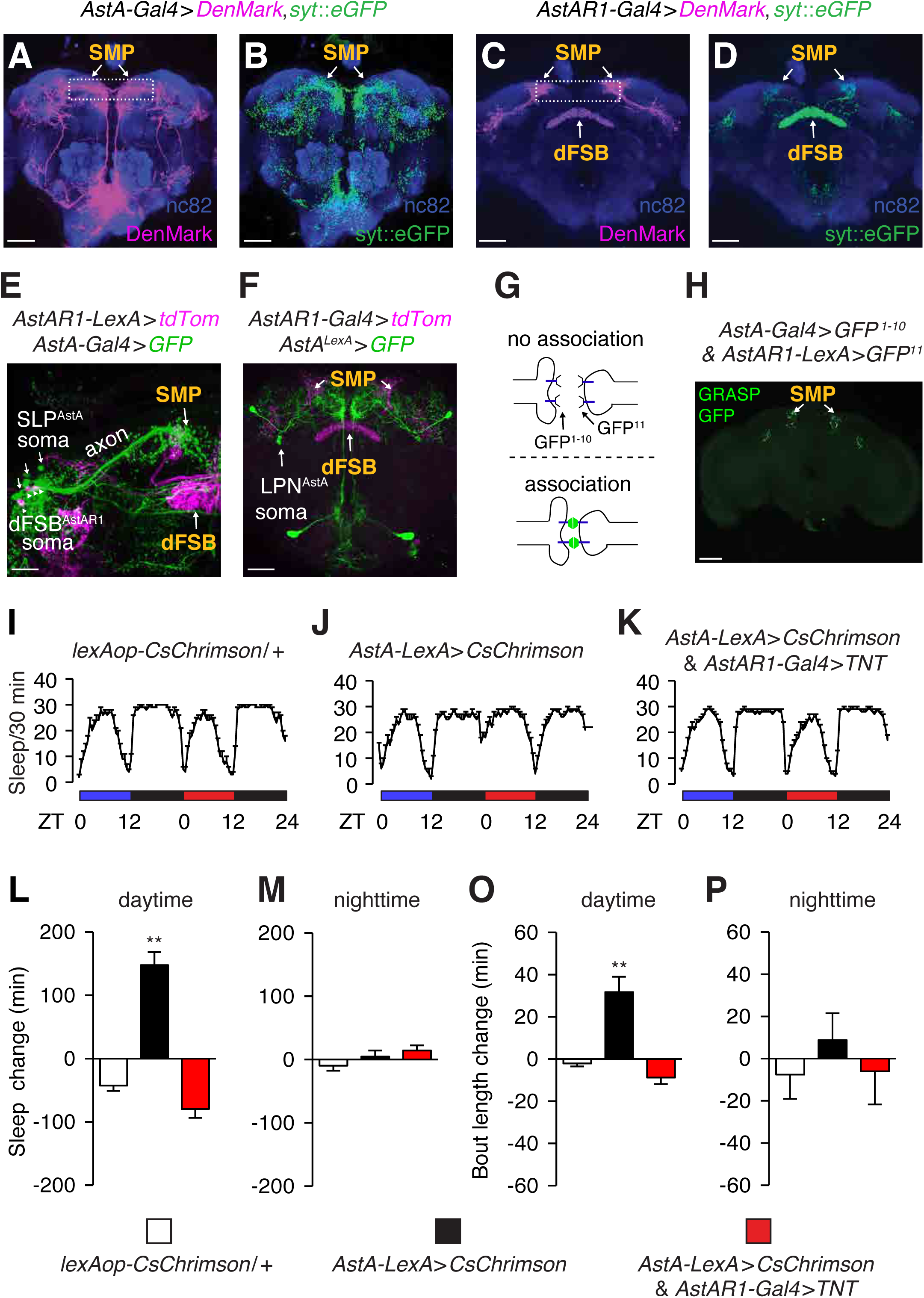
Anatomical and functional connectivity between *AstA-Gal4* positive and dFSB^AstAR1^ sleep-promoting neurons. (**A — B**) Immunostaining of whole-mount brains from flies expressing *UAS-DenMark* and *UAS-syt::eGFP* under the control of the *AstA-Gal4* (*AstA-Gal4>DenMark>syt::eGFP*). (A) Anti-dsRed (recognizes DenMark), magenta. The box indicates the SMP region. (B) Anti-GFP (recognizes syt::eGFP), green. The scale bars represent 40 *μ*m. (**C — D**) Immunostaining of whole-mount brains from *AstAR1-Gal4>DenMark >syt-eGFP* flies. (C) Anti-dsRed (recognizes DenMark), magenta. The box indicates the SMP region. (D) Anti-GFP (recognizes syt::eGFP), green. The scale bars represent 40 *μ*m. (**E**) Immunostaining of a whole-mount brain from a fly expressing the *AstAR1* reporter (*AstAR1-LexA>tdTomatd*) and the *AstA* reporter (*AstA-Gal4>GFP*). Both reporters showed innervation in the SMP region of the brain (Figure S2). The scale bar represents 20 *μ*m (**F**) Immunostaining of whole-mount brains of the *AstAR1* reporter (*AstAR1-Gal4>mCherry*) and the *AstA* reporter (*AstA^lexA^>GFP*). Both reporters stained the SMP region of the brain. The scale bar represents 40 *μ*m. (**G**) Cartoon illustrating the GRASP assay. Fluorescence is produced only when the two segments of GFP associate on the extracellular surfaces of adjacent cells. (**H**) Image of GRASP GFP fluorescence revealing close association between *AstA-Gal4* and *AstAR1-LexA* neurons in the brain. The scale bar represents 60 *μ*m. (**I — K**) Sleep profiles of the indicated flies under 12 hour blue light/dark and 12 hour red light/dark cycles. (**L — M**) Quantification of the changes in daytime and nighttime sleep (in minutes) due to neuronal activation by red lights in flies expressing CsChrimson. The changes in total sleep and the average bout lengths were calculated by subtracting these sleep parameter values obtained during the blue-light/dark cycles from those obtained during the red-light/dark cycles. (**O — P**) Quantification the change in daytime and nighttime sleep bout length due to neuronal activation by red lights in flies expressing CsChrimson. n=15−31. Error bars, SEMs. **p<0.01, one-way ANOVA with Dunnett’s test.

As a second approach to address whether the AstA (SLP^AstA^ and LPN^AstA^) and AstAR1 (dFSB^AstAR1^)-positive neurons form close associations, we performed double-labeling experiments. We examined the relative positions of the neuronal projections labeled by the *AstA* and *AstAR1* reporters, and found that the SMP region was innervated by both SLP^AstA^ neurons and dFSB^AstAR1^ neurons (Figure 4E and S2B). We also found that LPN^AstA^ neurons (*AstA^lexA^*) and dFSB^AstAR1^ neurons (*AstAR1-Gal4*) innervated a common SMP region (Figures 4F and S2).

We then used GFP Reconstitution Across Synaptic Partners (GRASP) (Feinberg et al., 2008; Gordon and Scott, 2009) to probe for close associations between *AstA* (LPN^AstA^ and SLP^AstA^) and dFSB^AstAR1^ neurons that could be due to synaptic connections between these two sets of sleep-promoting neurons. In this approach, two non-functional fragments of mCD4-GFP (mCD4-GFP^1–10^ and mCD4-GFP^11^) are expressed on the extracellular surfaces of different cells. GFP fluorescence is reconstituted only if the two fragments are brought into contact (Figure 4G). Consistent with close associations, we detected GFP fluorescence in the SMP region of the brain, where LPN^AstA^, SLP^AstA^ neurons (expressing *AstA-Gal4>UAS-mCD4-GFP^1–10^*) and dFSB^AstAR1^ neurons (expressing *AstAR1-LexA>lexAop-mCD4-GFP^11^*) send their projections (Figure 4H).

### dFSB^AstAR1^ neurons function downstream of SLP^AstA^ and LPN^AstA^ neurons to promote sleep

To test whether or not synaptic transmission is required for the sleep-promoting effect of SLP^AstA^ and LPN^AstA^ neurons, we expressed *UAS-trpA1* under control of the *AstA-Gal4* in the presence or absence of a temperature-sensitive dynamin (*UAS-shi^ts^*), which blocks synaptic transmission at the non-permissive temperature (29°C). At this same temperature (29°C), TRPA1 is activated, thereby stimulating the neurons. In the absence of *shi^ts^*, thermo-stimulation of the neurons with TRPA1 promotes sleep (Figure S4J). However, blocking neurotransmission with *shi^ts^* at t29°C prevents the sleep-promoting effect due to neuronal activation by TRPA1 (Figure S4I).

Close anatomical association between LPN^AstA^ and SLP^AstA^ and dFSB^AstAR1^ neurons suggests possible synaptic transmission between them. We therefore investigated if dFSB^AstAR1^ neurons function downstream of LPN^AstA^ and SLP^AstA^ neurons to promote sleep. We expressed *lexAop-CsChrimson* using an *AstA-LexA* driver that contains the same DNA regulatory sequences as the *AstA-Gal4*, and found that neuronal activation with red light promotes an increase in daytime sleep (Figures 4I, 4J, 4L and 4M), and consolidates daytime sleep into longer bouts (Figure 4O). However, CsChrimson activation of LPN^AstA^ and SLP^AstA^ neurons did not promote sleep or lead to bout consolidation if we blocked neurotransmission of dFSB^AstAR1^ neurons with tetanus toxin (TNT; Figures 4K—P).

### Glutamate is the sleep-promoting neurotransmitter used by LPN^AstA^ and SLP^AstA^ neurons to activate dFSB^AstAR1^ neurons

To identify the neurotransmitter released by the LPN^AstA^ and SLP^AstA^ neurons to enhance sleep, we used RNAi to knockdown genes essential for the synthesis or packaging of neurotransmitters. We found that the increase in sleep due to constitutive activation of these neurons by NaChBac (*AstA-Gal4* and *UAS-NaChBac*) was reduced by RNAi directed against the gene encoding the vesicular glutamate transporter (VGLUT; Figures 5A and 5F) (Daniels et al., 2006). In contrast, the elevation in sleep induced by NaChBac was not diminished when we performed RNAi on genes required for the synthesis of other major neurotransmitters, including acetylcholine (*ChAT*), γ-aminobutyric acid (GABA; *GAD1*) and biogenic amines (*VMAT*) (Figures 5B—5F). These observations suggest an important role for glutamate in conferring the sleep-promoting effect of LPN^AstA^ and SLP^AstA^ neurons.

**Figure 5.**
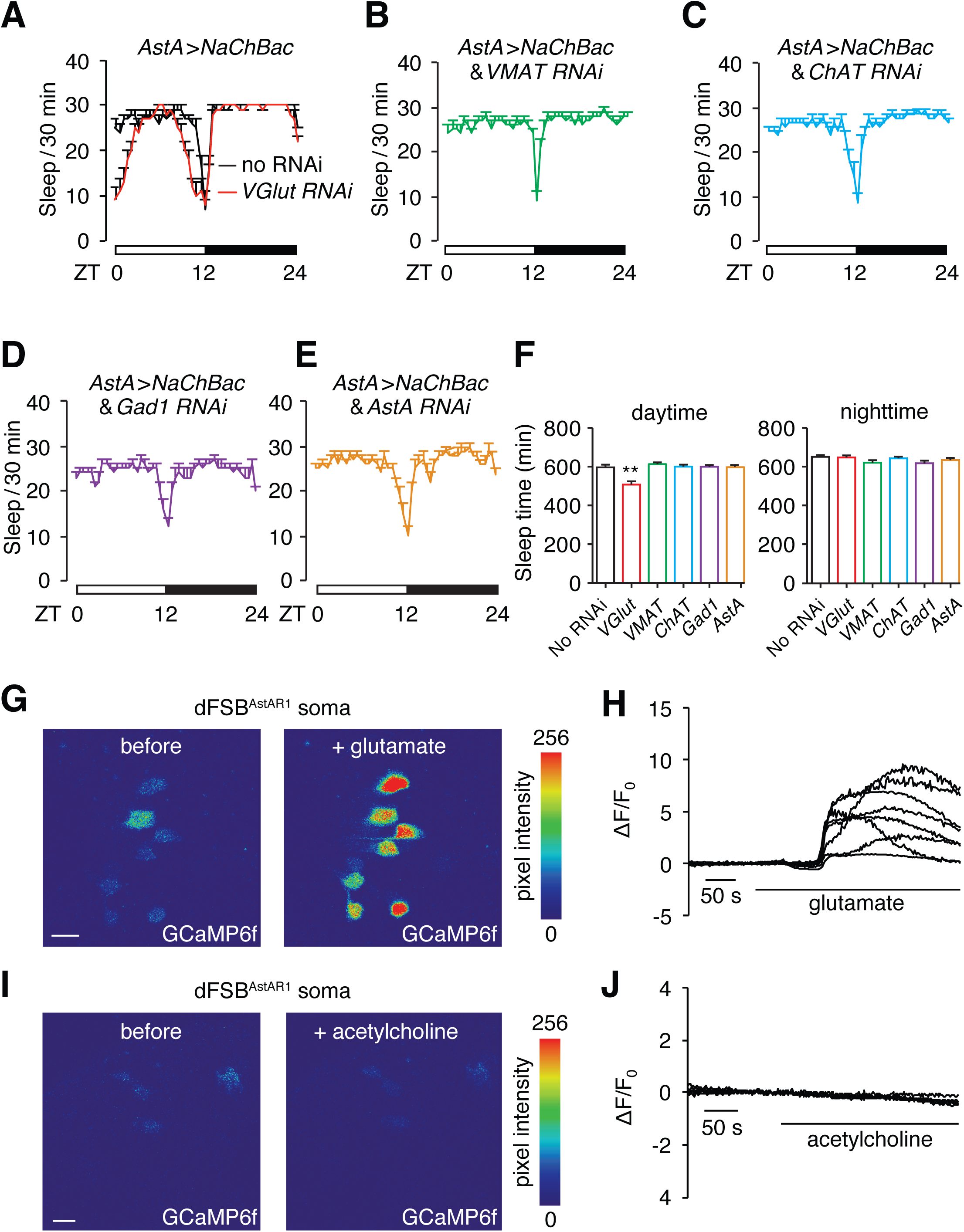
Glutamate is the excitatory neurotransmitter in *AstA-Gal4* positive neurons, which activates dFSB^AstAR1^ neurons and promotes sleep. (**A**) Sleep profile of flies expressing *UAS-NaChBac* under control of the *AstA-Gal4* (*AstA>NachBac*) with or without *VGlut* RNAi (*UAS-VGlut^RNAi^*). (**B—E**) Sleep profile of *AstA>NaChBac* flies expressing *VMAT-RNAi, ChAT-RNAi Gad1-RNAi*, or the *AstA-RNAi* transgenes. (**F**) Quantification of daytime sleep and nighttime sleep displayed by the indicated flies. n=29−55. Error bars, SEMs. **p<0.01, one-way ANOVA with Dunnett’s test. (**G**) Representative images of GCaMP6f fluorescence in a fly expressing *UAS-GCaMP6f* under the control of the *AstAR1-Gal4* (*AstAR1>GCaMP6*). Shown are the soma of dFSB^AstAR1^ neurons before and after bath application of 50 mM glutamate. (**H**) Representative traces showing changes in GCaMP6f fluorescence (ΔF/F_0_) in *AstAR1>GCaMP6* flies upon bath application of 50 mM glutamate. (**I**) Representative images of GCaMP6f fluorescence in *AstAR1>GCaMP6* flies before the after bath application of 50 mM acetylcholine. (**J**) Representative traces showing changes in GCaMP6 fluorescence (ΔF/F_0_) in *AstAR1>GCaMP6* flies upon bath application of 50 mM acetylcholine. The scale bars in G and I represents 20 *μ*m.

To test whether the dFSB^AstAR1^ neurons respond to glutamate, we employed an *ex-vivo* brain preparation so that we can image neuronal activity in freshly dissected brains. We expressed a fluorescent Ca^2+^ sensor (*UAS-GCaMP6f*) using the *AstAR1-Gal4* and imaged the soma of dFSB^AstAR1^ neurons after applying glutamate to the bath. We found that addition of glutamate caused an increase in GCaMP6f fluorescence (Figures 5G and 5H). In contrast, these neurons did not respond to acetylcholine—another major excitatory neurotransmitter in the *Drosophila* brain (Figures 5I and 5J).

### GABA is required in dFSB^AstAR1^ neurons to promote sleep

To screen for the neurotransmitter synthesized in dFSB^AstAR1^ neurons that is essential for stimulating sleep, we silenced expression of genes required for neurotransmitter synthesis or packaging. Thermoactivation of dFSB^AstAR1^ neurons with TRPA1 (*AstAR1>trpA1*) enhances sleep (Figures 6A, 6B, 6D and 6E). However, this effect was eliminated when we used RNAi to knockdown *Gad1* in dFSB neurons (*AstAR1>trpA1* and *Gad1* RNAi; Figures 6C and 6F). In contrast, we still observed significant sleep-promoting effects resulting from TRPA1-induced activation of dFSB^AstAR1^ neurons when we used RNAi to suppress production of other major neurotransmitters (Figures 6G—6I). These results indicate that GABA is a critical sleep-enhancing neurotransmitter synthesized in dFSB^AstAR1^ neurons.

**Figure 6.**
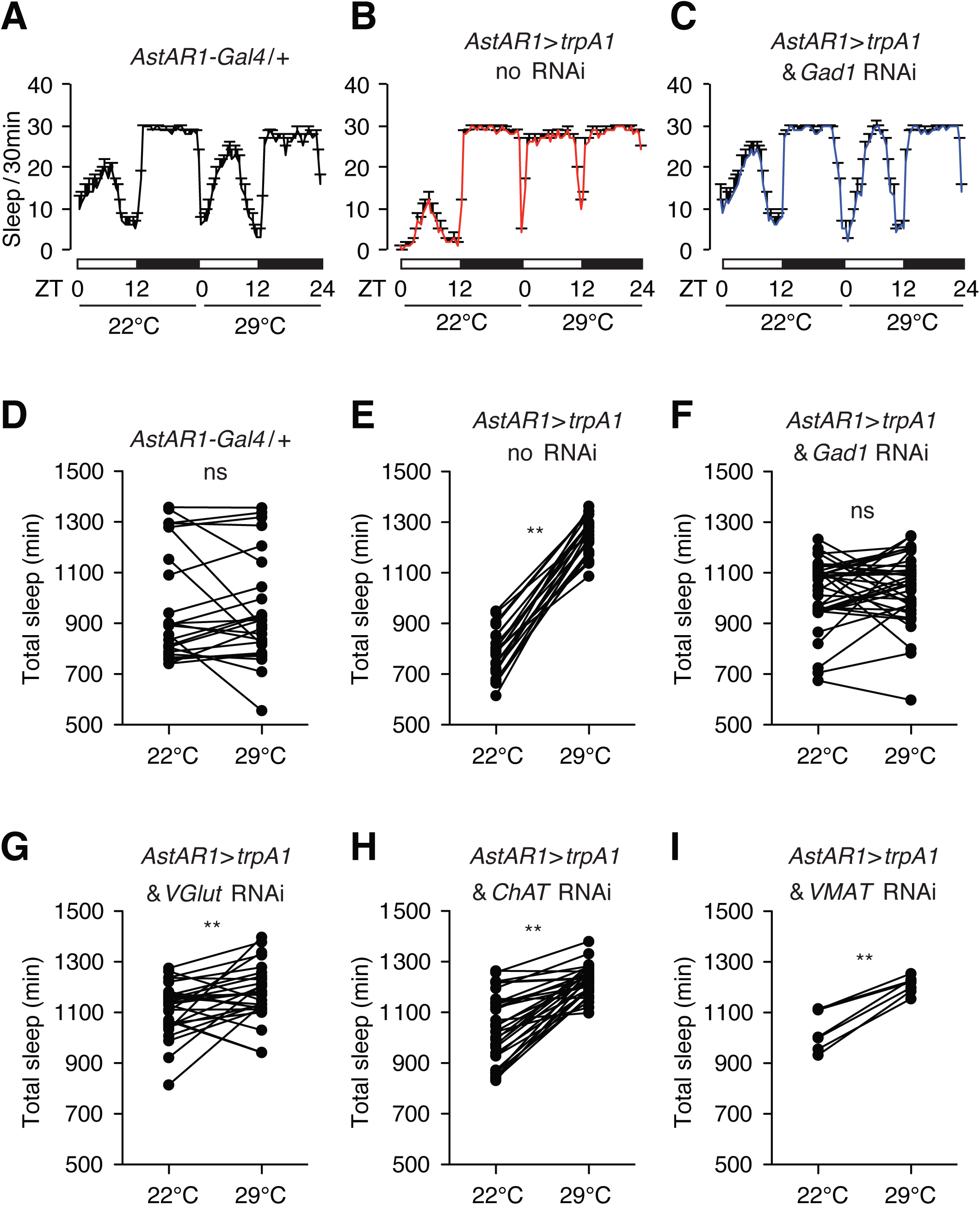
GABA is required in *AstAR1-Gal4* neurons to promote sleep. Testing for suppression of the increased sleep in *AstAR1>trpA1* (*AstAR1-Gal4* in combination with *UAS-trpA1*) flies, after RNAi-mediated knockdown of genes required for synthesis of neurotransmitters. We performed sleep profiles at 22°C (no TRPA1 activation) and 29°C (activation of TRPA1) as indicated. (**A — C**) Sleep profiles of control flies (*AstAR1-Gal4/+*) and *AstAR1>trpA1* flies with or without *Gad1* RNAi. (**D — I**) Quantification of total sleep-time for one 24 hour light/dark cycle at 22°C and one 24 hour light/dark 29°C of the indicated flies. **p<0.01, paired Student’s t-test. n=7−37.

### dFSB^AstAR1^ sleep-promoting neurons inhibit arousal-promoting neurons

Octopamine (OA) appears to increase arousal, as nighttime sleep is reduced upon feeding flies OA (Crocker and Sehgal, 2008) or by activating tyrosine decarboxylase 2 (TDC2)-expressing neurons with NaChBac (*tdc2-Gal4>NaChBac*) (Crocker et al., 2010) (Figure S5A). Nighttime sleep in *tdc2-Gal4>NaChBac* flies is fragmented, as the number of sleep bouts increases dramatically and the average length of each nighttime sleep bout is reduced significantly (Figures S5B and S5C).

We wondered whether dFSB^AstAR1^ neurons might down-regulate the activity of TDC2 positive arousal (OAA) neurons. We used two approaches to test for possible close associations between these two sets of neurons, which would be required for synaptic connections. First, we performed double-labeling experiments. The *Tdc2* reporter labeled neurons that project to multiple layers of the FSB, including the dorsal layer where dFSB^AstAR1^ neurons also project (Figure S5D). We then expressed dendritic (DenMark) and axonal (syt::eGFP) markers under control of the *tdc2-Gal4* to visualize the potential input and output sites of the Tdc2-positive (OA) neurons. Most of the OA neuronal projections in the dFSB were axonal (syt::eGFP; Figure S5E). In addition, there was a single dendritic layer in the dFSB labeled with DenMark (Figure S5E). To determine whether dFSB and OA neurons were in close contact, we performed a second approach—GRASP analysis. We observed GRASP signals in multiple areas of the brain. Of note, the OA neurons, which express the *Tdc2-Gal4*, form close connections with the dFSB^AstAR1^ sleep-promoting neurons (Figure S5F).

To determine if activation of dFSB^AstAR1^ neurons inhibited octopaminergic arousal (OAA) neurons, we expressed the ATP-activated cation channel P2X2 in dFSB^AstAR1^ neurons, and imaged OAA neuronal activity with GCaMP3 (Yao et al., 2012). After applying ATP, we observed a dramatic reduction in GCaMP3 fluorescence (Figures S5G—S5H). These data indicate that dFSB^AstAR1^ sleep-promoting neurons inhibit OAA neurons (Figure S2B).

### Inhibition of dFSB^AstAR1^ sleep-promoting neurons by dopaminergic arousal neurons

A dopaminergic arousal circuit projects to the dFSB (Liu et al., 2012; Ueno et al., 2012b), raising the possibility that dopaminergic arousal (DAA) neurons negatively regulate dFSB^AstAR1^ sleep-promoting neurons. We analyzed the anatomical relationship of dFSB^AstAR^ and DAA neurons by labeling the dFSB^AstAR^ neurons with GFP (*AstAR1-Gal4>GFP*) and the DAA neurons with antibodies that recognize TH (tyrosine hydroxylase), an enzyme required for dopamine biosynthesis. Consistent with the proposal that DAA neurons regulate dFSB^AstAR1^ neurons, anti-TH and anti-GFP labeled neighboring neurons in the dFSB (Figure S6A).

To image the relative innervation patterns of the two classes of neurons, we labeled DAA neurons with a GFP reporter (*TH-Gal4>GFP*) that is expressed in a majority of neurons that stain with anti-TH (Figures S6B and S6C) (Friggi-Grelin et al., 2003; Liu et al., 2012). In addition, we labeled the dFSB^AstAR1^ neurons with tdTomato (*AstAR1-LexA>tdTomato;* Figures 7A and 7B). We found that the DAA neurons project to two layers within the dFSB: the dorsal FSB layer (dFSB) and the ventral FSB layer (vFSB; Figure 7B). The dorsal layer overlaps with the projections from dFSB^AstAR1^ neurons (Figure 7B), suggesting possible synaptic connection between DAA and dFSB^AstAR1^ neurons. We used GRASP to assess whether the DAA and dFSB^AstAR1^ neuronal membranes were in close proximity—a prerequisite for forming synaptic connections. We expressed *UAS-mCD4-GFP^1–10^* and *lexAop-mCD4-GFP^11^* under the control of the *TH-Gal4* and *AstAR1-LexA*, respectively, and observed GRASP signals in regions of the brain containing the receptive field of the dFSB neuronal projections (Figures 7C and S6D). We did not detect GFP signals in brains expressing only one of the two GFP fragments (Figures S6E and S6F).

**Figure 7.**
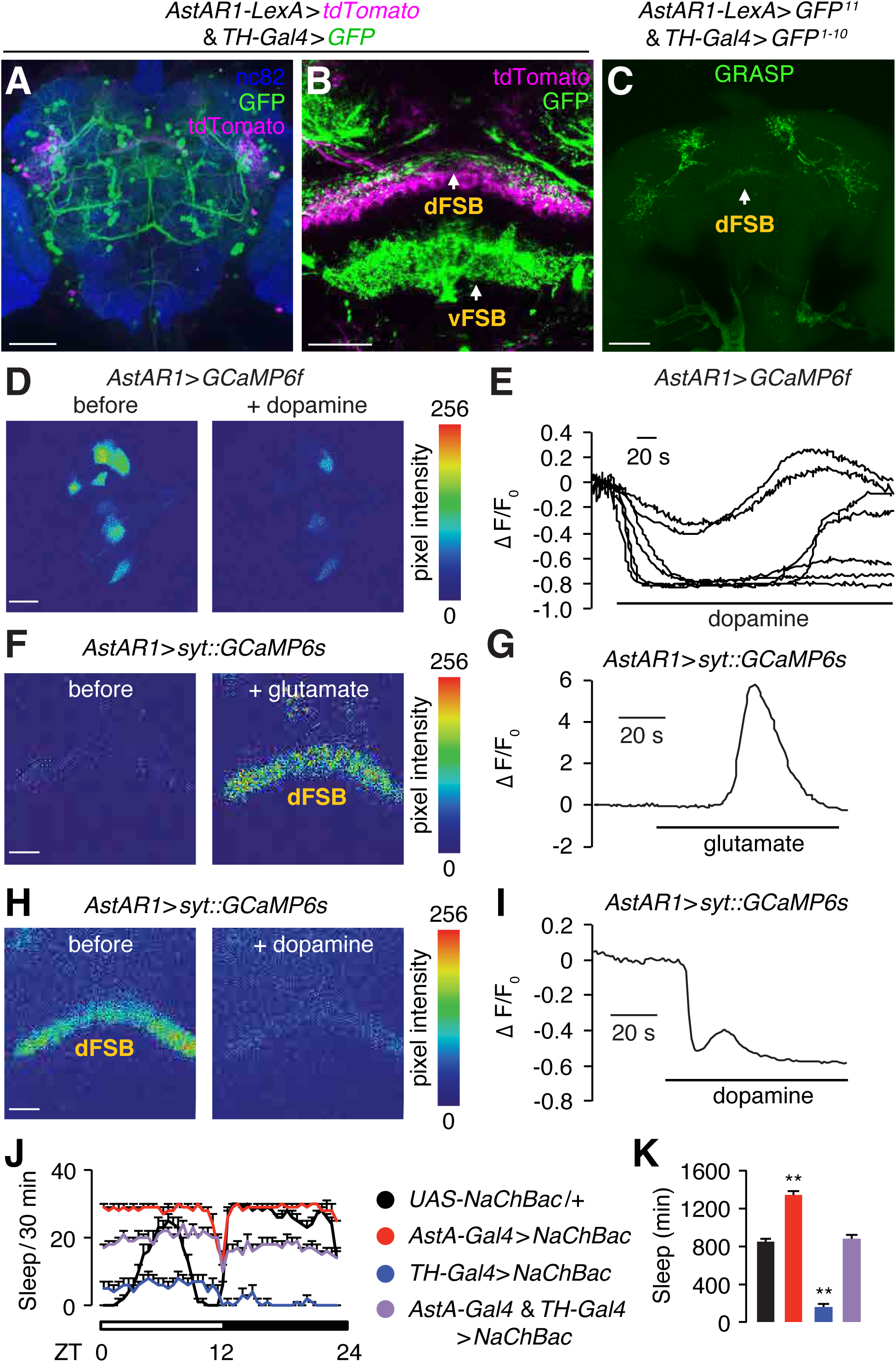
Dopaminergic neurons inhibit the activity of sleep-promoting dFSB^AstAR1^ neurons. (**A, B**) Immunostaining of brains expressing a dopaminergic reporter (*TH-Gal4>GFP*, stained with anti-GFP, green), and a reporter labeling dFSB^AstAR1^ sleep-promoting neurons (*AstAR1-LexA>tdTomato*, stained with anti-DsRed to label tdTomato, magenta). Anti-nc82 in A, blue. The dFSB and vFSB regions are indicated in B. The scale bars represents 60 and 20 *μ*m in A and B, respectively. (**C**) GRASP GFP fluorescence revealing close association between sleep-promoting dFSB^AstAR1^ neurons and dopaminergic neurons in the brain. The scale bar represents 60 *μ*m. (**D**) Representative image of GCaMP6f fluorescence in the cell soma of a brain from *AstAR1>GCaMP6f* flies before and during bath application of 10 mM dopamine. The scale bars in D, F and H represent 20 *μ*m. (**E**) Representative traces of changes in GCaMP6f fluorescence (ΔF/F_0_) in *AstAR1>GCaMP6f* flies upon bath application of 10 mM dopamine. n=7. (**F**) Representative image of syt::GCaMP6s fluorescence in axonal terminals from a brain from *AstAR1>syt::GCaMP6s* flies before and during bath application of 50 mM glutamate. (**G**) Representative trace showing the change in syt::GCaMP6s fluorescence (ΔF/F_0_) in *AstAR1>syt::GCaMP6s* axonal terminals upon bath application of 50 mM glutamate. (**H**) Representative images of syt::GCaMP6s fluorescence in in *AstAR1>syt::GCaMP6s* axonal terminals before and during bath application of 10 mM dopamine. (**I**) Representative trace showing the change in syt::GCaMP6s fluorescence (ΔF/F_0_) in *AstAR1>syt::GCaMP6s* axonal terminals upon bath application of 10 mM dopamine. (**J**) Activation of dopaminergic neurons using NaChBac antagonizes the sleep-promoting effect of *AstA-Gal4* neuronal activation. Shown are the sleep profiles of the indicated genotypes. (**K**) Quantification of total sleep time of one combined light/dark cycle using flies of the indicated genotypes. n=12−24. Error bars, SEMs. *p<0.05, **p<0.01 one-way ANOVA with Dunnett’s test.

To address whether dopamine inhibits sleep-promoting dFSB^AstAR^ neurons, we expressed *UAS-GCaMP6f* using the *AstAR1-Gal4* and imaged the soma of the neurons with projections that extend to the dFSB. After applying dopamine to the buffer, there was a large reduction in GCaMP6f fluorescence (Figures 7D and 7E), indicating that dopamine inhibits sleep-promoting dFSB^AstAR1^ neurons.

To test whether or not activation of DAA neurons inhibits dFSB^AstAR1^ neurons, we expressed the ATP-activated cation channel P2X2 in DA neurons (*TH-Gal4>P2X2*) and monitored changes in the activity of dFSB^AstAR1^ neurons with GCaMP3 (*AstAR1-LexA>GCaMP3*). After applying ATP, we observed a significant reduction (p<0.01, Unpaired Student’s t-test) in GCaMP3 fluorescence, relative to the effect of applying the imaging buffer only (Figures S7A and S7B). In contrast, when we activated dFSB^AstAR1^ neurons with ATP (*AstA-Gal4>P2X2*), we observed a 2.2 ±0.2 fold increase in GCaMP3 fluorescence in dFSB^AstAR1^ neurons (Figures S7C and S7D).

### Effect of simultaneous activation of dFSB^AstAR1^ and DAA neurons

Sleep-promoting dFSB^AstAR1^ neurons receive inputs from *AstA-Gal4* positive neurons (LPN^AstA^ and SLP^AstA^) that project to the SMP region (Figure S2). The outputs from the dFSB^AstAR1^ neurons are transmitted through their axonal projections to the dFSB. The GRASP data described above (Figures 4H) indicate that glutamatergic inputs from *AstA-Gal4* neurons are restricted primarily to the SMP, and are not detected in the dFSB. However, DAA neurons form connections at multiple projection areas of the sleep-promoting dFSB^AstAR1^ neurons, including the SMP and dFSB (Figures 7C and S2B).

We wondered if dopamine suppresses Ca^2+^ dynamics of dFSB^AstAR1^ neurons at their axonal terminals located in the dFSB region, thereby negatively regulating their output. To test this idea, we expressed syt::GCaMP6s (Cohn et al., 2015), in dFSB^AstAR1^ neurons, and confirmed its localization at its terminals in the dFSB by staining with anti-GFP (Figure S7E). We then imaged Ca^2+^ dynamics at the axonal terminals of dFSB^AstAR1^ neurons using an *ex-vivo* brain preparation. Consistent with the ability of glutamate to activate these neurons, application of glutamate to the bath solution resulted in a large elevation of Ca^2+^ (Figures 7F and 7G). When we applied dopamine to the bath, we observed a reduction in Ca^2+^ levels in the axonal terminals of the dFSB^AstAR1^ neurons (Figures 7H and 7I). Applying either of two other biogenic amines (octopamine and tyramine) did not change the Ca^2+^ dynamics in dFSB^AstAR1^ neurons (Figure S7F).

Since LPN^AstA^ and SLP^AstA^ provide positive inputs to dFSB^AstAR1^ sleep-promoting neurons, while DAA neurons are inhibitory to these neurons (Figure S2B), we tested the effects on sleep when these two groups of neurons were activated simultaneously. Activating DAA neurons (*TH-Gal4>NaChBac*) lead to a drastic reduction in total sleep time, relative to the control (*UAS-NaChBac/+;* 158 ±37 min versus 849 ±32 min; Figures,7J and 7K). As described above, activating *AstA-Gal4* neurons (LPN^AstA^ and SLP^AstA^) promotes sleep (1343 ±36 min; Figures 7J and 7K). Of significance, co-activating DAA and LPN^AstA^/SLP^AstA^ neurons with NaChBac lead to a level of total sleep time (881±46 min) similar to the control flies (849±32 min; Figures 7J and 7K).

To characterize the consequences of co-activating DAA and LPN^AstA^/SLP^AstA^ neurons in greater detail, we analyzed two features of sleep patterns: sleep bout number and sleep bout length. When we used the *AstA-Gal4* only to express *UAS-NaChBac* (*AstA>NaChBac*), sleep bouts were significantly lengthened (Figure S7G). Conversely, activating DAA neurons (*TH>NaChBac*) resulted in fragmented sleep, with a significantly reduced sleep bout length, which was most obvious for nighttime sleep (Figure S7G). When LPN^AstA^/SLP^AstA^ and DAA neurons were activated simultaneously with the *AstA-Gal4* and *TH-Gal4*, respectively, the sleep bouts were shorter than exhibited by the control, but were comparable to flies in which only the DAA neurons were activated (*TH>NaChBac;* Figure S7G).

Additionally, activating both LPN^AstA^/SLP^AstA^ and DAA neurons caused the flies to initiate many more sleep episodes, as indicated by a significant increase in the sleep bout number (Figure S7H).

We then compared the episodes of wakefulness of these flies. In control flies (*UAS-NaChBac*), wake bout length is much longer during the day than at night (daytime, 48.9 ±5.1 min.; nighttime, 9.3 ±1.1 min; Figures S7I). Activating sleep-promoting LPN^AstA^/SLP^AstA^ neurons (*AstA-Gal4>NaChBac*) lead to a reduction of the bout length of daytime wakefulness, while activating arousal-promoting DAA neurons increased the bout length of nighttime wakefulness (Figures S7I). Remarkably, when the two groups of neurons were activated simultaneously, both daytime and nighttime activities were fragmented (Figures S7I).

## Discussion

A regular sleep pattern is achieved through coordinated temporal segregation of sleep-promoting and arousal-promoting neuronal activity. However, the underlying neuronal circuits that are necessary to achieve synchronization between these opposing types of neuronal inputs are poorly understood. Nevertheless, neurons projecting to the dFSB of the central complex are proposed to be the effector arm of the sleep homeostat (Donlea et al., 2014; Liu et al., 2016). In our study, we identified multiple sleep and arousal neurons that comprise a neuronal network that enables flies to maintain a consolidated sleep pattern.

We identified sleep-promoting neurons that use glutamate as their neurotransmitter. Glutamate receptors are proposed to promote sleep in *Drosophila* (Tomita et al., 2015). Yet, the specific synaptic sites of sleep-promoting glutaminergic neurotransmission was unclear. Our results pinpoint one such type of sleep-promoting synapse. Using the GRASP technique, we found that axons extending from LPN and SLP neurons form close associations with dendrites of dFSB sleep-promoting neurons in the SMP region (Figure S2). Moreover, we found that the LPN and SLP neurons express the neuropeptide, Allatostatin-A (AstA), which has been linked to sleep promotion (Hentze et al., 2015), while the dFSB neurons express the reporter for the *Allatostatin-A Receptor 1* (*AstAR1*). Using *ex-vivo* Ca^2+^ imaging experiments, we found that glutamate released from LPN^AstA^ and SLP^AstA^ neurons acts as an excitatory neurotransmitter to activate dFSB^AstAR1^ neurons.

Dopamine is an arousal neuromodulator in both *Drosophila* and mammals (Andretic et al., 2008; Herrera-Solís et al., 2017; Liu et al., 2012; Ueno et al., 2012a) and in flies can switch dFSB neurons from an active to a quiescent state (Pimentel et al., 2016). Our GRASP analyses reveals that dopaminergic arousal (DAA) neurons form close associations at various locations along the dendritic and axonal regions of dFSB^AstAR1^ neurons. These include the SMP region where dFSB^AstAR1^ neurons receive glutaminergic input (Figure S2B). In addition, based on staining of dendritic and axonal makers, we propose that DAA neurons form synapses with axons of dFSB^AstAR1^ neurons in the dFSB region (Figure S2B). In support of this latter conclusion, we performed Ca^2+^ imaging and found that dopamine down-regulates Ca^2+^ levels at the axonal terminals of dFSB^AstAR1^ neurons. Thus, we propose that dopamine also contributes to arousal by suppressing the synaptic output of sleep promoting dFSB^AstAR1^ neurons.

dFSB^AstAR1^ neurons receive both sleep-promoting glutaminergic and arousal-promoting dopaminergic inputs. We found that simultaneous activation of these opposing inputs causes the flies to rapidly shift between sleep and wake states. Consequently, the flies are unable to experience long sleep bouts at night or stay awake for long periods during the day. Our observations argue for a requirement of temporal segregation of the input of sleep-promoting SLP^AstA^ /LPN^AstA^ neurons and the TH-positive DAA neurons, which regulate dFSB^AstAR1^ neurons in an opposing fashion.

An additional question is the mechanism through which dFSB^AstAR1^ neurons, which make up the effector arm of the *Drosophila* sleep-homeostatic circuit (Donlea et al., 2014; Liu et al., 2016), promote sleep. GABA is an inhibitory neurotransmitter, and has been linked to sleep promotion in flies and mammals (Agosto et al., 2008; Chung et al., 2017; Jones, 2017). We demonstrate that dFSB^AstAR1^ neurons use GABA as their sleep-promoting neurotransmitter. Moreover, we found that key targets for GABA are octopaminergic arousal-promoting (OAA) neurons (Figure S2). Based on GRASP analysis, we found that dFSB^AstAR1^ neurons and OAA neurons are in close association, and may form synapses. Moreover, activating dFSB^AstAR1^ neurons inhibits OAA neurons. This inhibitory effect, which is mediated by GABA, might serve to maintain long sleep bouts at night, since activating OAA neurons leads to fragmentation of nighttime sleep. Given that the dFSB region is part of the central complex, which integrates different types of sensory information (Wolff et al., 2015), we suggest that the GABA released by dFSB^AstAR1^ neurons provides inhibitory modulation that increases the threshold for sensory responses in sleeping animals.

## AUTHOR CONTRIBUTIONS

J.N. and C.M designed the experiments, interpreted the data and wrote the manuscript with the input from T.O. J.N. performed most of the experiments. T.O. contributed to the Ca^2+^ imaging and immunohistochemistry. A.E. and A.S.G. performed immunohistochemistry and behavior assays. H.H and A.V. contributed to the initial behavioral screen (Figure S1), and characterization of the reporter expression patterns identified in the screen.

## ACKNOWLEDGMENTS

The work was supported by a grant to C.M. from the National Institute on Deafness and other Communication Disorders (DC007864) and the National Institute of Allergy and Infectious Disease (DP1AI124453). We thank Drs. Orie Shafer and Julie Simpson for transgenic flies, Drs. Mark Wu and Sha Liu for guidance on performing sleep assays and Dr. William Joiner for sharing software. We also thank the Bloomington Stock Center for providing fly stocks.

## Star✶Methods

### Key Resources Table

| Primary Antibodies | Source | Identifier |
| --- | --- | --- |
| Rabbit anti-dsRed | Clontech | Cat. 432486 |
| Rabbit anti-GFP | ThermoFisher Scientific | Cat. A11122 |
| Chicken anti-GFP | ThermoFisher Scientific | Cat. A10262 |
| Mouse anti-GFP | Sigma-Aldrich | Cat. G6539 |
| Mouse anti-Brp (nc82) | DSHB | Antibody Registry ID: AB 2314866 |
| Rat anti-Timeless | Amita Sehgal lab | Jang *et al.* PLoS Genet. 2015 |
| Mouse anti-AstA | DSHB | Antibody Registry ID: AB 528076 |
| Rabbit anti-TH | EMD Millipore | Cat. AB152 |

| Secondary Antibodies | Source | Identifier |
| --- | --- | --- |
| Goat anti-mouse IgG1 | ThermoFisher Scientific | Alexa Fluor® 488, 568, 647 |
| Goat anti-rabbit IgG | ThermoFisher Scientific | Alexa Fluor® 488, 568 |
| Goat anti-rat IgG | ThermoFisher Scientific | Alexa Fluor® 555 |
| Goat anti-chicken | ThermoFisher Scientific | Alexa Fluor® 488 |

| Other chemicals | Source | Identifier |
| --- | --- | --- |
| Mounting media | Vector Laboratories | Cat. H-1000 |
| Triton X-100 | Sigma-Aldrich | Cat. T8787 |
| L-glutamic Acid | Sigma-Aldrich | Cat. G1251 |
| Dopamine | Sigma-Aldrich | Cat. H8502 |
| ATP | Roche | Cat. 10127531001 |
| Octopamine | Sigma-Aldrich | Cat. O0250 |
| Tyramine | Sigma-Aldrich | Cat. T2879 |

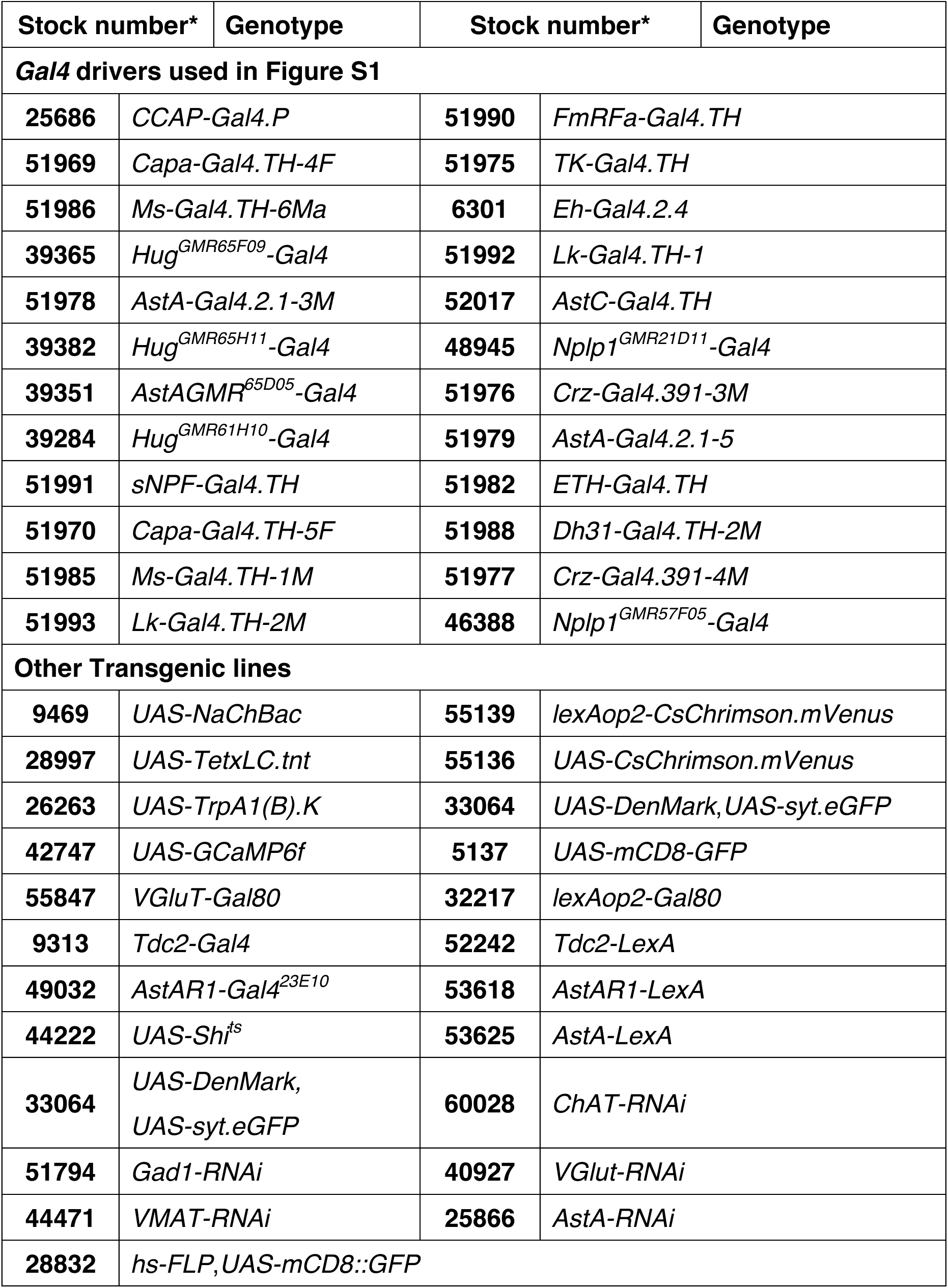

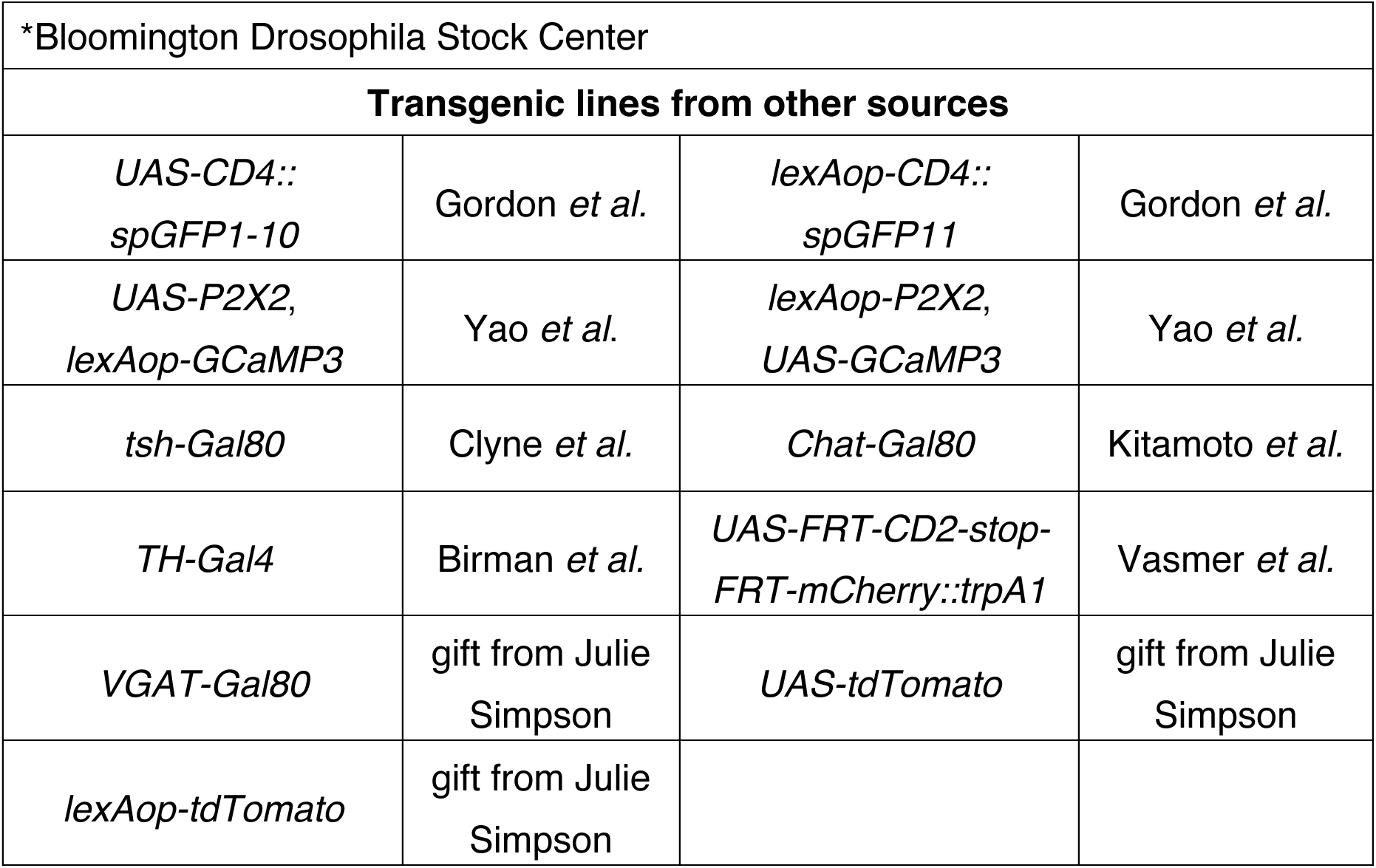

### Contact for Reagent and Resource Sharing

Further information and requests for reagents should be directed to and will be fulfilled by the Lead Contact, Craig Montell.

### Star Methods

#### Animals

Flies (*Drosophila melanogaster*) were cultured on cornmeal-agar-molasses medium under 12 hour light/dark cycle at room-temperature and ambient humidity. Detailed information regarding specific stains and genotypes is provided in the Key Resources Table section.

#### Sleep Behavior

To measure fly sleep, we used 3–7 day old female flies. Individual flies were loaded into glass tubes (provided with the Drosophila Activity Monitoring (DAM) system (TriKinetics Inc.). The tubes each contained 5% sucrose and 1% agarose as the food source at one end and a small cotton plug at the other end. Flies were entrained for ≥2 days before activity data were collected for analysis. The activity data were collected in one-minute bins for further processing using MATLAB (MathWorks)-based software (Koh et al., 2008). If no activity was detected by the DAM system for more than 5 minutes, the fly was considered to be in a sleep status. Sleep assays were performed under white LED lights at 25°C unless otherwise mentioned. For optogenetic stimulation, flies were cultured in regular food supplied with *all-trans* retinal in the dark for 24 hours before loaded into the glass tubes to perform activity measurements.

#### Screening for Peptidergic Neurons and *AstA1R1* Neurons that Affect Sleep

To identify peptidergic neurons that affect sleep, we used a panel of *Gal4* lines (listed in the Key Resource Table) to drive expression of *UAS-trpA1*. The putative peptidergic neurons labeled by these lines were activated using thermogenetics as described in the following section (Activation of Neurons using Optogenetics and Thermogenetics), and the effects on sleep were determined using 8–24 female F1 offspring per assay.

To identify *AstAR1* neurons that function downstream of *AstA-Gal4* neurons (SLP^AstA^ and LPN^AstA^), we obtained 25 *Gal4* lines from Janelia Farm (Jenett et al., 2012) that contained DNA from the putative *AstAR1* enhancer/promoter regions, and used these lines to drive expression of *UAS-NaChBac* (Nitabach et al., 2006). We used 8–16 female F1 flies to perform the initial sleep assays. The negative control was *UAS-NaChBac* only. The *23E10-Gal4* line, which is expressed primarily in the dFSB (Jenett et al., 2012), displayed the most pronounced sleep-promoting effect and phenocopied the sleep-promoting effect of the *AstA-Gal4* lines. We repeated the experiments with the *23E10-Gal4* line after the initial screen two additional times.

#### Activation of Neurons using Optogenetics and Thermogenetics

To measure fly sleep using optogenetics, we used 3–7 day-old female flies. Flies were cultured in regular food supplied with *all-trans* retinal in the dark for 24 hours before loading individual flies into the glass tubes provided with the DAM assay platform provided by Trikinetics Inc. The tubes contained 5% sucrose and 1% agarose as the food source at one end and a small cotton plug at the other end. Flies were entrained for 3 days under blue LED lights. On the 4^th^ day, the LED lights were switched to red. Activity data were collected in one-minute bins for further processing using Sleep-Lab software. The effects on sleep as a result of optogenetic activation of neurons were quantified as the sleep change under red lights on day 4 verses blue lights on day 3.

We used two thermogenetic approaches in combination with the *Gal4/UAS* system to manipulate neuronal activity: 1) we expressed TRPA1, and activated the channel and the neurons at 29°C (Parisky et al., 2008), and 2) we expressed a temperature sensitive allele of Shibire (Shi^ts^) (Kitamoto, 2001; Kosaka and Ikeda, 1983) and silenced the neurons after shifting the temperature to 29°C. To conduct these analyses, we introduced 3–7 day old female flies that were cultured on standard into Trikinetics glass tubes with 5% sucrose + 1% agarose as the food source at one end and a small cotton plug at the other end. Flies were entrained for 3 days under white LED lights at 22°C and the temperature was switched to 29°C on the day 4. Activity data were collect in one-minute bins for further process using Sleep-Lab software (Joiner et al., 2013). The effects on sleep as a result of thermogenetic manipulation of neuronal activity were quantified as the sleep change (in minutes) at 29°C (4^th^ day) verses 22°C (3^rd^ day).

#### Whole-mount Brain Immunohistochemistry

For whole-mount brain immunohistochemistry, brains were dissected in phosphate-buffered saline and 0.3% Triton X-100 (PBST). The brains were fixed with 4% paraformaldehyde (PFA) in PBST at room temperature for 20 minutes. Brains were briefly washed two times in PBST, blocked with PBST and 5% normal goat serum (NGS) at 4°C for 1 hour, and incubated overnight with primary antibodies diluted in PBST and 5% NGS at the following dilutions: 1) (1:1000) chicken anti-GFP, rabbit anti-dsRed, rabbit anti-TH antibody, and rat anti-Timeless, 2) (1:500) rabbit anti-GFP, 3) (1:200) nc82, 4) mouse anti-GFP antibody (1:100), and 5) (1:50) mouse anti-AstA. After three washes in PBST, we incubated the brains overnight in secondary antibodies corresponding to the primary antibodies used. Secondary antibodies are diluted 1:1000 in PBST and 5% NGS (refer to the Key Resource Table for the list of secondary antibodies). The brains were washed three times in PBST, and mounted in VECTASHIELD (Vector Laboratories) on glass slides. Images were acquired with an upright Zeiss LSM 700 confocal miscroscope using 20X, 40X (oil) or 63X (oil) lenses.

#### Mosaic Expression of *mCherry::trpA1* using the *FlpOut* Method

To use mCherry::TRPA1 to activate different subsets of neurons that express the *AstA-Gal4*, we conducted a mosaic analysis to express *mCherry::trpA1* in random subsets of *AstA-Gal4* neurons. To do so, we combined *hs-Flp* and *AstA-Gal4* transgenes with the *UAS-mCD8::GPF* transgene, and the *UAS-FRT-CD2-stop-FRT-mCherry::trpA1* transgene (Vasmer et al., 2014). The animals were maintained at room temperature (~22°C). To remove the *stop* cassette, we induced *hs-FLP* expression by heat shocking 3^rd^ instar larval at 37°C for 1 hour. The animals were then returned to room temperature (~22°C) after the heat shock. We refer to these animals as *AstA>FlpOut-mCherry::trpA1* flies. We used 3–7 day-old female *AstA>FlpOut-mCherry::trpA1* flies to conduct the sleep measurements following the thermogenetic procedure described above. To tabulate the percentage of flies with a given amount of sleep change due to thermal activation of mCherry::TRPA1, we used the following calculation: sleep at 29°C minus sleep at 22°C.

To image the neurons, we dissected the brains of *AstA>FlpOut-mCherry::trpA1* flies that showed either sleep-promoting effects (sleep change > 100 minutes) and that did not show sleep-promoting effects (sleep change < 100 minutes). We then stained the brains with anti-GFP and anti-dsRed as described above (Whole-mount Brain Immunohistochemistry).

#### GRASP Analysis

To conduct the GRASP analyses (Feinberg et al., 2008; Gordon and Scott, 2009), we expressed *lexAop-CD4::spGFP1* and *UAS-CD4::spGFP1–101* under control of the *AstAR1-LexA* and either the *AstA-Gal4* or the *TH-Gal4*. To image GRASP signals, the brains were dissected in PBST, fixed with 4% PFA in PBST at room temperature for 20 minutes, and briefly washed with PBST before mounting in VECTASHIELD (Vector Laboratories) and viewing the fluorescence. To enhance GRASP signals, we performed immunostaining using mouse anti-GFP.

#### Ca^2+^ Imaging in Brains

To perform Ca^2+^ imaging on whole-mount brains, expressing GCaMP3, GCaMP6f or syt::GCaMP6s, the flies were anaesthetized on ice and the brains were dissected into AHL buffer (108 mM NaCl, 5 mM KCl, 8.2 mM MgCl_2_, 2 mM CaCl_2_, 4 mM NaHCO_3_, 1 mM NaH_2_PO_4_, 5 mM trehalose, 5 mM sucrose, and 5 mM HEPES pH 7.5), and transferred to an imaging chamber with a glass bottom, which was made from cover slides. The brains were immobilized with a metal harp and imaged using an inverted Zeiss LSM 800 confocal microscope. The regions of interest in the soma and neuronal processes were scanned in the time-lapse mode using 4–6 Z stacks. 30 or more cycles (60–90 seconds total) were imaged for checking the stability of the sample (not sliding and floating) in the imaging chamber prior to adding chemicals to the bath. The average intensity during the last 10 of these 30 pre-cycles (before adding the chemicals) was used as the baseline (F) to calculate ΔF.

#### Quantification and Statistical Analysis

To analyze sleep behavior, multiple animals were tested and the sleep index of individual animals were quantified independently. The numbers of animals tested were all biological replicates with numbers (n) indicated in all figure legends. We used Student’s t-tests to compare two group of samples. In most cases the tests were unpaired. We used paired Student’s t-tests for comparing the effects of thermogenetic activation with TRPA1 in Figure 6. For comparing multiple groups, we used One-way ANOVA followed by the Dunnett test. Error bars indicates SEMs with indicated p values for statistical analysis. * Indicates p<0.05 and ** indicates p<0.01.

